# Viscoelastic characterization of cells in microfluidic channels with 3D hydrodynamic focusing

**DOI:** 10.1101/2025.04.17.649258

**Authors:** Benedikt Hartmann, Felix Reichel, Conrad Möckel, Jochen Guck

## Abstract

The viscoelastic nature of biological cells has emerged as an increasingly important research subject due to its relevance for cellular functions under physiological and pathological conditions. Advancements in microfluidics have made this technology a promising tool to study the viscoelasticity of cells. However, significant challenges remain, including the complex distribution of stresses acting on cells depending on the channel geometry, and the difficulty of keeping cells in the focal plane for imaging. Here, we report a new approach using hyperbolic channels for measuring cell viscoelasticity. A channel height much larger than the typical cell size minimized shear stresses so that normal stresses in the hyperbolic region dominated the stress distribution. Reducing the complexity of the stress-strain relationship allowed us to use polyacrylamide microgel beads to calibrate the stress curve. Additionally, we introduced 3D hydrodynamic focusing which enabled us to focus cells and microgel beads in the center of the channel. Finally, Kelvin-Voigt and power-law rheology models were employed to extract the mechanical properties of microgel beads and human leukemia HL60 cells. The measurement technique described here will help establish the viscoelastic properties of cells as an important readout in biophysical research in health and disease.

## Introduction

Viscoelasticity of cells plays a key role in numerous biological processes, e.g. cell migration^1,2^, immune response^3–5^, and other physiological and pathological cell functions^6–9^. Until now, numerous studies showed that cell stiffness and pathological changes are correlated, yet the causal relationships are still unknown. Understanding their origin, pathways, and interplay with other factors, such as gene expression, holds the potential for utilizing mechanical properties as label-free biomarkers for diagnosis and poses a potential therapeutic target.^10–12^ As transcriptional profiles become increasingly available, we are one step closer to identifying specific genes that determine a cell’s mechanical phenotype, leading to the ultimate goal of being able to control mechanical properties.^13,14^ Reliable methods to assess the mechanical properties of cells are therefore a necessity.

Several techniques have been developed to quantify cell viscoelasticity. Established methods, such as atomic force microscopy^15^, micropipette aspiration^16^, or optical stretching^17^ usually suffer from low throughput and require tedious operation.

On the other hand, microfluidic approaches are becoming more widely adopted.^18–22^ These methods leverage various channel geometries, probing cells across a wide range of spatial and temporal scales. Recently, Reichel *et al*.^23^ introduced a microfluidic method that utilizes a hyperbolic channel design to probe the mechanical properties of cells. However, their method requires a careful estimation of shear and tensile stresses, due to significant shear contributions from the velocity profile across the channel height. In addition, investigations on the rheological properties of the carrier medium in shear and extensional flow are required to understand the evolution of the stresses along the channel axis. Taken together, these points require specialized technical knowledge and thus represent an obstacle to the widespread establishment and application of the method.

In this study, we present a new technique based on an advanced hyperbolic channel design. By using a channel height of 90 µm, we achieved minimal shear stress contributions from the velocity profile across the channel height and simplified the stress distribution. This helped to avoid different modes of deformation, as extensional flow causes a tensile stretching of the cells into an ellipse-like shape while shear stresses cause a bullet-like shape.^24^ Additionally, we employed 3D hydrodynamic focusing presented by Zhao *et al*.^25^ to keep cells in the center plane of the channel, thus improving imaging and throughput.

Stresses in the channel were characterized using Polyacrylamide (PAAm) microgel beads, which react instantaneously to changes in stress due to their nearly perfect elastic behavior.^26,27^ We also employed PAAm microgel beads to validate our approach by showing that we can recover expected mechanical properties as measured with real-time deformability cytometry (RT-DC).^28^ Furthermore, we showed that our method can detect changes in the mechanical properties of HL60 cells treated with latrunculin B, a drug inhibiting actin polymerization.

Our results illustrate how a combination of experimental techniques can be used to optimize the viscoelastic characterization of cells using hyperbolic channels. This method has a great potential to contribute to the field of biomechanics as it could serve, for example, as a screening platform to advance the discovery of drugs altering cellular mechanical properties or mechanics-related pathways.^13,29^

## Methods

### Microfluidic chip design and fabrication

The channel design is depicted in Figure 1 and is similar to the chip design introduced by Reichel *et al*.^23^ The height of the channel was *H* = 91 ± 4 µm. Next to the sample inlet are two additional inlets on either side for the vertical focusing flows. After a second, horizontal focusing step using sheath flows, the sample flows through a 2.2 mm long and *w*_*u*_ = 500 µm wide channel before it reaches the *L*_*c*_ = 300 μm long hyperbolic contraction. The shape of the hyperbolic contraction is described by the following relation^23^

**Figure 1.**
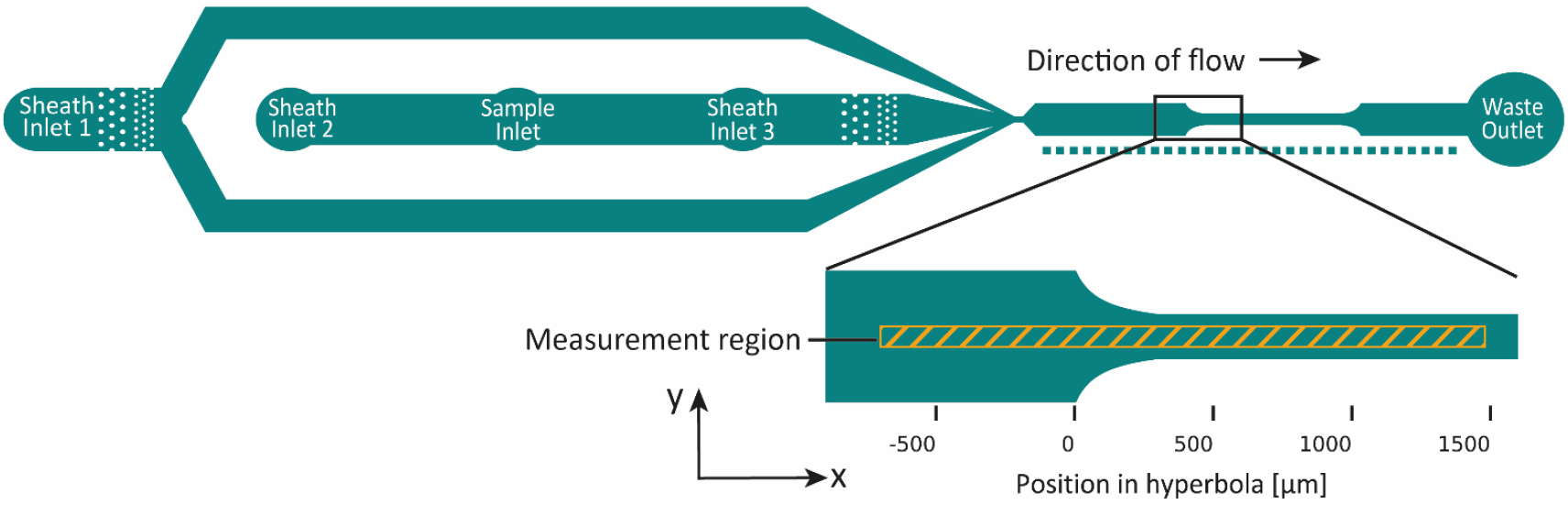
Overview of the channel design. From the left: Sheath and sample inlets, focusing region, hyperbolic channel, outlet. Horizontal focusing is provided by sheath flow from the sides via inlet 1. Vertical focusing is achieved via sheath inlets 2 and 3 next to the sample inlet. The measurement region comprises the hatched orange area shown in the inset and was 54 µm wide and, in total, 2180 µm long.

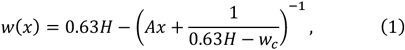

with

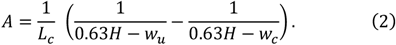

The channel width after the hyperbola is *w*_*c*_ = 160 µm and continues straight for 2 mm before opening again with a mirrored hyperbolic part into a 500 µm wide channel. At the end of that channel is the waste outlet. The microfluidic chips were produced using standard soft lithography. Briefly, the mold was produced by applying a 90 µm thick layer of SU8-3050 photoresist (MicroChemical GmbH, Germany) on a 4-inch silicon wafer using a spin coater (WS-650Mz-23NPP, Laurell, USA) at 500 RPM for 10 seconds followed by 2000 RPM for 30 seconds. After a waiting period of 10 minutes, the wafer with the layer of photoresist was soft-baked for 25 minutes at 95 °C. A chromium mask with the chip design was illuminated by UV light with an energy density of 220 mJ/cm^2^ (MA6 Gen4, Suess MicroTec GmbH, Germany) to transfer the design onto the photoresist, followed by baking at 65 °C for 1 minute and 95 °C for 5 minutes. Final development of the mold was done with mr-DEV600 for 5 minutes and 15 seconds followed by isopropanol treatment for 30 seconds.Polydimethylsiloxane (PDMS, VWR 634165S, SYLGARD 184) was prepared with a base-to-curing agent ratio of 10:1 and poured onto the mold, degassed, and cured for 75 minutes at 78 °C. The replica of the mold was then cut into individual chips, inlet holes punched with a 1.5 mm biopsy puncher (pfm medical AG, Germany) and bonded to a glass cover slip (40 mm × 24 mm, thickness 2, Hecht, Germany) using air plasma treatment (Plasma system Atto, Diener Electronic, Germany) at 75 W for 2 minutes, with 20 sccm air flow at 0.4 mbar pressure.

### Polyacrylamide microgel beads (PAAm beads)

All microgel beads used in this study were produced as described by Girardo *et al*.^26^ In total, five different batches were used, three for the stress characterization and two for validation. Beads sizes and Young’s moduli were measured using RT-DC^28,30^ and are given together with the total monomer concentration in the supplementary material (Table S1, Fig. S1, Fig. S2).

### Cell culture and drug treatment

HL60/S4 cells (ATCC Cat# CRL-3306, RRID:CV-CL_II77) were cultured in RPMI 1640 medium (Thermo Fisher #A1049101) with 10% heat-inactivated fetal bovine serum in an incubator at 37 °C at 5% CO_2_. Cell density was kept between 10^5^ cells/mL and 10^6^ cells/mL and subcultured every 48–72 hours. Cells were seeded on the day before the experiment at a concentration of 10^5^ cells/mL per flask. On the day of an experiment, the cells were counted and the viability checked to be >90%. Treatment with latrunculin B (latB, latrunculia magnifica, EMD Millipore Corp., USA, product number 428020, LOT 3541421, CAS # 76343-94-7) was performed as follows: A stock solution of 10 mM latB was prepared by dissolving the powder in Dimethyl Sulfoxide (DMSO, VWR Chemicals, USA, Catalog # N182, CAS # 67-68-5). In a next step, this stock was diluted further with DMSO to get 10 000 times the desired latB concentration. This ensured an equivalent DMSO concentration of 0.01 v/v% in all experiments. Finally, the latB solution was diluted by a factor of 10 000 in the carrier medium to reach final concentrations of 0 nM and 100 nM latB.

### Sample preparation

#### PAAm microgel beads

Beads were taken from the Eppendorf tube in which they were stored and where they formed a pellet at the bottom. For an experiment, 2 µL of beads were resuspended in 50 µL of carrier medium.

#### Cells

For experiments, cells were collected from the cell culture flasks, counted and their viability checked with Erythrosin B Stain (Logos Biosystems, Catalog # L13002). Experiments were performed only if the viability was >90 %. Cells were stored in an incubator during waiting periods. For an experiment, cells were centrifuged for 2 minutes at 200*g* and resuspended at a concentration of 5 × 10^6^ cells/mL in a final volume of 50 µL of the carrier medium containing the respective latB concentration. The samples were stored in an incubator (37 °C, 5% CO_2_) for 30 minutes and afterwards aspirated into the sample tubing with a negative flow rate of −2 µL/s to perform measurements.

### Carrier medium

A stock of the carrier medium was prepared by dissolving 2 g of methyl cellulose powder (4000 cPs, Alfa Aesar, 36718.22, CAS 9004-67-5) in 198 g of phosphate buffered saline (1X DPBS, Gibco Dulbecco, catalog # 14190144), resulting in a concentration of 1 w/w%. Before adding the powder, the osmolality of the DPBS was checked to be 280 – 290 mmol/kg (VAPRO Vapor Pressure Osmometer 5600, Wescor, USA). The pH was measured (Orion Star A211 benchtop pH meter, Thermo Scientific) and adjusted if needed by adding NaOH until a value of 7.4 was reached. Finally, it was filtered through a polyethersulfone (PES) filter, pore size 0.22 µm (Millex-GP, product number SLGP033RS, Merck Millipore, Ireland). The stock solution of the carrier medium was stored at 4 °C and taken out of the fridge before an experiment to equilibrate with room temperature.

### Experimental procedure

Experiments were carried out on an AcCellerator (Zellmechanik Dresden, Germany), a device for RT-DC that consists of an inverted microscope (Axio Observer Z1, Zeiss, Germany), syringe pumps (neMESYS low pressure syringe pump, Cetoni, Germany), and a high-speed camera (EoSens CL, MC1362, Mikrotron, Germany). Experiments were performed at 24 ± 1 °C. Imaging was performed using an objective lens with 20x magnification (Plan-Apochromat, x20/0.8 NA, product number 420650-9902, Zeiss, Germany). The resulting pixel size was 0.68 µm/px. The data was recorded using the acquisition software of the AcCellerator device, Shape-In 2. The frame rate was chosen depending on the flow rate to capture at least 60 images per object. The region of interest (ROI) had a size of 80 px × 1000 px or 54 µm × 680 µm.Four syringe pumps were used for the experiments. One glass syringe with 100 µL total volume (SETonic GmbH, Germany) was used for the sample, two 1 mL glass syringes (SETonic GmbH, Germany) were used for vertical sheath flow, and one 1 mL glass syringe was used for horizontal sheath flow. Syringes and microfluidic chip were connected by FEP tubing (1/16” OD, 0.03” ID, product number 1520XL, Postnova Analytics, Germany). Table 1 shows an overview of the total flow rates used and the corresponding flow rates for each syringe. All samples used in this study were measured at these flow rates.

**Table 1.** Overview of flow rates and frame rates.

| Total flow rate [μL/s] | Frame rate [fps] | Sample flow rate [μL/s] | Vertical sheath flow rate (per syringe) [μL/s] | Horizontal sheath flow rate [μL/s] |
| --- | --- | --- | --- | --- |
| 0.025 | 200 | 0.001 | 0.002625 | 0.01875 |
| 0.050 | 600 | 0.002 | 0.005250 | 0.03750 |
| 0.100 | 1200 | 0.002 | 0.011500 | 0.07500 |
| 0.200 | 1600 | 0.004 | 0.023000 | 0.15000 |

All syringes and their respective tubing were filled with carrier medium. The syringe for horizontal sheath flow additionally contained SiO_2_ microparticles (product number 56798, Sigma Aldrich) in the following ratio: 5 µL from the homogeneous particle suspension per 1.2 mL of carrier medium. These particles were used to determine the top and bottom of the channel by focusing on very slowly moving or stationary particles and noting down the z-position of the objective lens. This was used to calculate the correct position of the objective lens to image the channel’s center plane.Once filled, all syringes except the sample syringe were connected to the microfluidic chip and the channel was filled with carrier medium. The sample was prepared and aspirated into the tubing with a flow rate of −2 µL/s. Once the sample was loaded into the tubing, the tubing was connected to the microfluidic chip, and the flow rates per syringe were set according to Table 1.

### Data acquisition

First, a calibration measurement > 500 µm before the start of the hyperbolic region was performed at a total flow rate of 0.025 µL/s. Calibration of the strain was required because the PAAm microgel beads exhibited a non-zero strain value in that part of the channel despite the absence of any stresses large enough to deform them. This strain showed a dependency on the position within the ROI. We attribute this phenomenon to optical distortions in the setup and the low contrast of the beads due to their refractive index being similar to that of the surrounding medium. Consequently, camera noise and small brightness fluctuations affect the contour in a way that leads to higher strain. We fitted each strain calibration curve per bead sample with a polynomial function of order 12 and corrected all further data of that respective sample with it. This effect was not observed for cells, likely due to the much higher contrast to the background. However, the cells exhibited a finite strain value 500 µm before the hyperbolic part due to not being perfectly round.After the calibration measurement, four measurements per flow rate were recorded with the ROI starting at *x* = −700 µm, *x* = −200 µm, *x* = 300 µm, and *x* = 800 µm relative to the start of the hyperbola. This creates a 180 µm overlap between two consecutive measurements to facilitate stitching together of the data. Recordings of cells were stopped either after reaching 60 000 events or after a measurement time of 4 minutes to avoid large differences in total measurement time.

### Data processing

For each sample, the four measurements along the channel were analyzed with a custom written Python script to track objects and to compute features needed for the analysis. The median radius per sample was determined from the data at the beginning of the measurement region, between *x* = −700 µm and *x* = −360 µm relative to the start of the hyperbola, where there was no flow-induced deformation.The strain feature was calculated for each event as

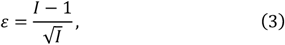

with the inertia ratio *I* of the event, i.e., the ratio of the second order central moments of the event contour along *x* and *y* direction and computed already by the acquisition software.^23^ This feature is more robust to noise impacting the event contour, in contrast to pixel-based features like the aspect ratio. The strain of the PAAm microgel beads was then corrected by the respective amount determined from the calibration measurement.The velocity of each tracked object was determined by the custom Python script and then a velocity curve over position in the channel was fitted with a generalized logistic function to get a smooth velocity curve for each sample. This function describes the asymptotic behavior of the velocity before and after the hyperbolic contraction and allows more flexibility than the logistic or sigmoid functions. The function is given by

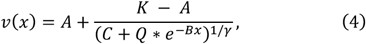

where *A* describes the left horizontal asymptote, *K* the right horizontal asymptote when *C* = 1, *C* rescales the right horizontal asymptote as *A* + (*K* − *A*)/(*C*^1/γ^) when *C* > 1, *Q* is related to the value *v*(0), *B* is the growth rate and γ affects the position of maximum growth.The characterization of the stress curves is explained in the results section. The full code used for the analysis is available online at http://hdl.handle.net/21.11101/0000-0007-FE52-F.

### Mechanical characterization

To calculate the mechanical properties of measured samples, we apply two common mechanical models, namely the Kelvin-Voigt model and power-law rheology. The constitutive equation for the Kelvin-Voigt model is given by

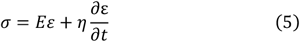

with one-dimensional stress σ, elastic modulus *E*, strain ε, and bulk viscosity η.For an arbitrary stress function, σ(*t*), the general solution of the Kelvin-Voigt is as follows^31,32^

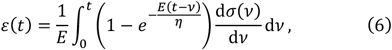

which can be further simplified by integration by parts and assuming that the objects are stress free in the wide channel before the hyperbola (σ(*t* = 0) = 0):

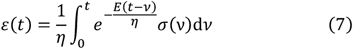

To consider the not perfectly round shape of the cells, we adjusted the formula by adding an additional parameter ε_offset_:

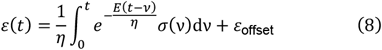

This formula is used to fit the recorded strain data to determine *E*, η, and from them, consequently calculate the relaxation time τ = η/*E*. The constitutive equation for the power-law is given by

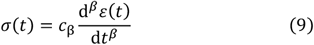

with the power-law exponent 0 < β < 1 and the material property *c*_β_.^31^ While β indicates a purely elastic material for β → 0 and a purely viscous behavior for β → 1, the interpretation of *c*_β_ is more difficult, as it has fractional units of [*c*_β_] = Pa s^β^. For β → 0, *c*_β_ can be interpreted as the stiffness of the material and in the case β → 1 as its viscosity.The general solution for the power-law reads^31^

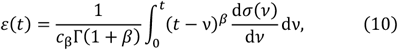

where Γ(…) denotes the Gamma function. Integration by parts and assuming that stresses are negligible before the hyperbola leads to

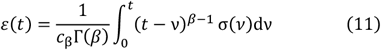

To consider the not perfectly round shape of the cells, we added again a parameter ε_offset_. The formula that was eventually used to determine power-law exponent β and the material property *c*_β_ reads

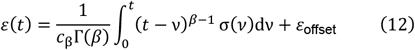

## Results and Discussion

### 3-D hydrodynamic focusing

3D-focusing of cells and cell-like objects has been achieved by combination of the established horizontal focusing^28^ and the vertical focusing presented by Zhao *et al*.^25^ The principle of vertical focusing is shown in Fig. 2A. Two sheath inlets provide the flow to force the sample into the vertical center plane of the channel. Due to the properties of laminar flow, the sample stays in this plane along the channel. This minimizes shear stresses from top and bottom channel walls and allows imaging in the same focal plane across the measurement region. Example pictures taken at different z-positions inside the channel are presented in Fig. 2B. In the image taken at *z* = 45 µm, the PAAm beads are barely visible because they are nearly perfectly in focus. The beads imaged at *z* = 59 µm, however, show a bright white halo around them, indicating that they are out of focus. Similarly, the beads imaged at *z* = 32 µm exhibit a dark border with a smaller, brighter inner halo as they are out of focus in the opposite direction compared to those at *z* = 59 µm.^33^ We achieved sufficient focusing to assume all objects are in the channel center where the shear stress is minimal. The flow rates of the sheath inlets could be increased to achieve even better focusing, especially horizontally, but increasing the sheath flow would also dilute the sample more, thereby reducing the throughput.

**Figure 2.**
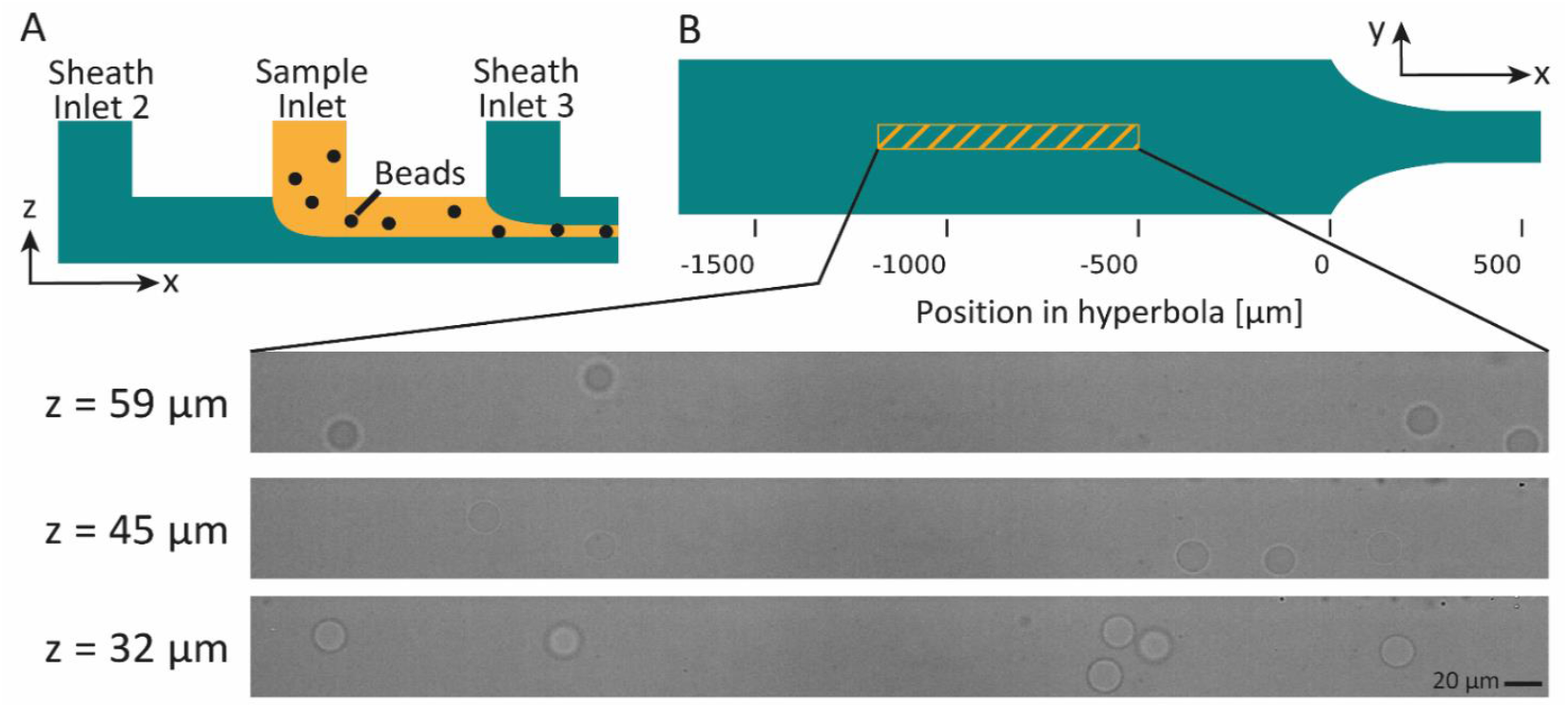
3D hydrodynamic focusing. A) Principle of vertical focusing: The cross-section of the channel is first filled with carrier medium from sheath inlet 2 before the sample is introduced via another inlet. This leads to a layer of liquid containing the sample at the top of the channel. The flow from sheath inlet 3 effectively pushes this layer down to the vertical center plane of the channel. The flow is laminar, so particles once focused in the inlet region stay in the correct plane further down the channel. B) Position of strain calibration measurement 500 µm before the start of the hyperbola. Images taken at different positions of the objective lens at *z* = 32 µm, *z* = 45 µm (channel center), and *z* = 59 µm illustrate this. For higher contrast, the grayscale value range of the images was set to 70 – 215 instead of 0 – 255. Calibration beads B (big + soft), measured at 0.2 µL/s flow rate.

In our experiments, we noticed impaired vertical focusing with microfluidic chips where the holes of the sample inlet or sheath inlets 2 and 3 were punched laterally off-center. In that case, the sample is pushed down unevenly depending on the position in the channel, and it would not even reach the center plane of the channel if the sample inlet is shifted to one side of the channel and sheath inlet 3 is shifted to the other. A reduced diameter of the sample inlet and precise punching of the inlet hole could further improve the lateral position of cells and their lateral spread in the inlet region, leading to more precise vertical focusing.

### Stress characterization using PAAm microgel beads

Next, we characterized the stress in the channel. For that purpose, we employed three different batches of PAAm beads as stress sensors^26^: Calibration samples A, B, and C, which differ in size and Young’s modulus (see Figs. S1 and S2 and Table S1). Although the carrier medium is a viscoelastic liquid with shear-thinning and non-zero normal-stress differences^34^, we assume that the PAAm beads are suitable to locally measure the apparent stress along the channel due to their almost purely elastic behavior.^26,27^ Their deformation will adjust almost immediately to stresses from their surrounding environment so that we can use it as a reasonable readout for the stress.Strain curves for all three samples were measured at the flow rates listed in Table 1. The apparent stress in the channel was calculated for each bead sample by multiplying the corrected strain with the Young’s modulus as measured by RT-DC (see Supplementary Materials and Methods and Fig. S2)

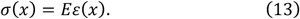

This is done under the assumption of linear elasticity, as the maximum strain for any of the calibration samples was below 0.15.^23^ Calibration samples A and B were measured in a channel of height *H* = 96 µm, calibration sample C was measured in a channel of height *H* = 88 µm. To compensate for different bead sizes and channel heights and therefore differences in shear stress, we rescaled the apparent stress using the confinement, κ ≡ diameter/channel height. The rescaled, apparent stress can therefore be written as

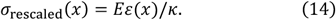

Exemplary data for the flow rate of 0.1μ*L*/*s* is shown in Fig. 3.

**Figure 3.**
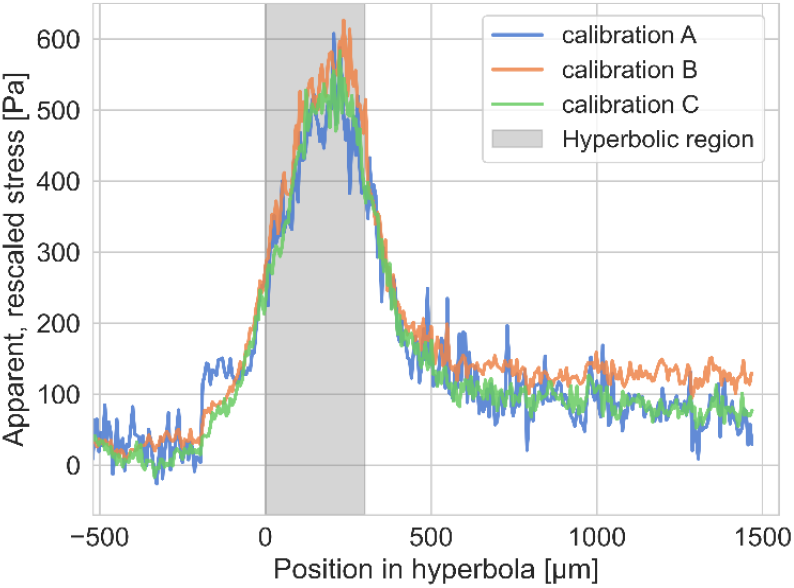
Individual curves of apparent, rescaled stress over position along the channel for three different calibration samples of PAAm beads. The total flow rate was 0.1 µL/s, over 4000 beads were measured for each sample. The data is binned, with each bin covering 6 µm. The three curves overlap well despite different sizes, Young’s moduli, and channel heights.

The apparent stress curves of all three calibration samples overlap well and show negligible stress before the hyperbolic region (marked in gray). The stress reaches a peak around the middle of the hyperbolic region before decreasing and settling on a constant value. This constant value is due to shear stresses, because the flow velocity is increased in the smaller channel cross section in this part of the chip. There is no extensional flow after the hyperbolic region. The stress curve of calibration sample A shows more fluctuations than the others. This is likely due to the fact that these are the stiffest of all calibration beads used, resulting in less deformation in the channel and consequently a lower signal-to-noise ratio.The position *x* = −360 µm relative to the start of the hyperbola was used as a reference point for the stress and strain curves in the further analysis for all four flow rates. The apparent stress curves of all three samples were combined, binned and the mean value per bin was calculated. These stress curves (of the four different flow rates) were used in the following sections for the characterization of the mechanical properties of two batches of PAAm beads and latB treated HL60 cells.

### Mechanical characterization of PAAm microgel beads

Two samples of PAAm beads, validation A and B, were measured to demonstrate the utility of our method for the mechanical characterization of microscopic particles. As for the calibration samples, a measurement 500 µm before the hyperbola was recorded to determine the strain correction.To calculate the mechanical properties of the validation samples, we applied the Kelvin-Voigt model and power-law rheology. The general stress curves determined in the previous section describe the apparent stress over the position in the channel. To apply the mechanical models (described in Eq. 7 and Eq. 11, respectively), which depend on time, *t*, the general stress curves, σ(*x*), and the strain curves of the samples, ε_A,B_(*x*), need to be transformed into stress and strain curves over time, σ(*t*) and ε_A,B_(*t*), respectively.For that purpose, we used the velocity over position curve, *v*(*x*), for each sample and flow rate to recalculate the time corresponding to the respective positions in the channel according to the following formula ^23^

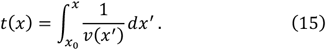

Fitting of the resulting data was performed using a custom Mathematica script. The data range was restricted to *x* ≥ −360 µ*m* relative to the beginning of the hyperbola, as there was no strain before that point. The stress curve was smoothened by applying the built-in function “GaussianFilter” and subsequently interpolated to obtain stress values at the exact time points of the strain data.The results of the Kelvin-Voigt model-based analysis are shown in Figs. 4A-C. The Young’s moduli are given in Fig. 4A and show comparable values for all four flow rates. For comparison, the median Young’s moduli measured with RT-DC are also shown in Fig. 4A as dashed lines, the shaded areas denote the standard deviations of the RT-DC measurements. The Young’s moduli determined with our method are lower than the expected values determined by RT-DC, but are within the margins of uncertainty.

**Figure 4.**
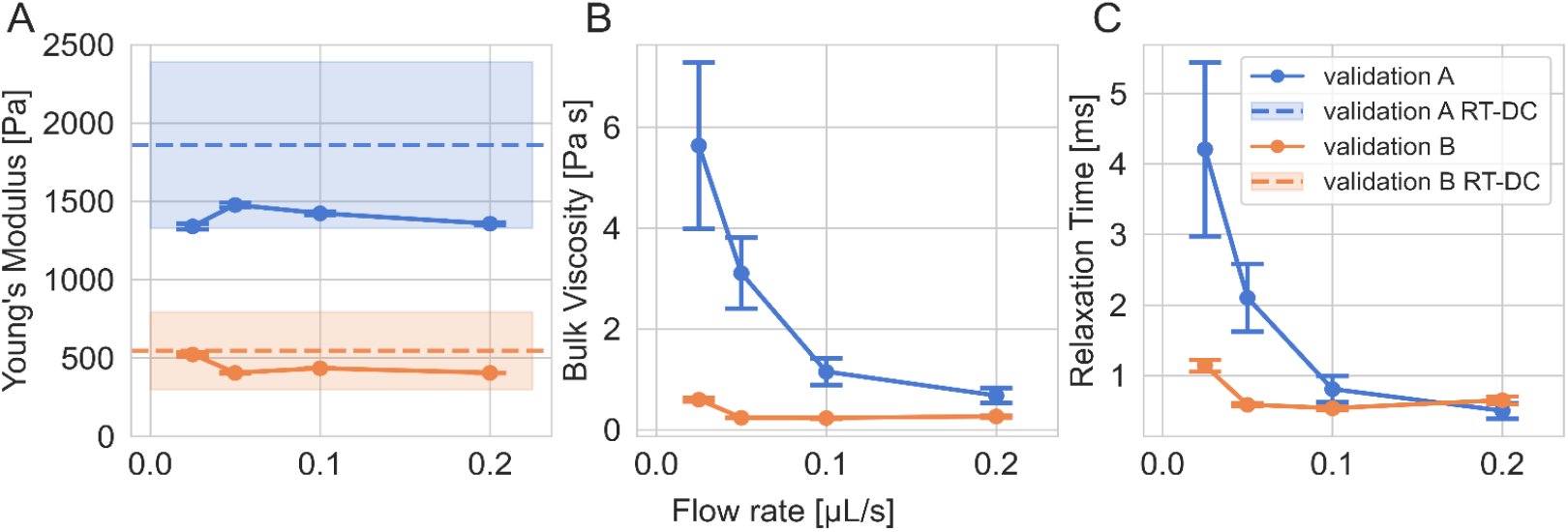
Mechanical characterization of PAAm microgel beads. A) Young’s modulus measured in the hyperbolic channel, error bars denote uncertainties from the fit. The dashed lines show the median results from RT-DC measurements for comparison, the shaded area denotes the standard deviation of the RT-DC data. B) Bulk viscosity of the PAAm microgel beads. C) Relaxation times of PAAm microgel beads.

Generally, it should be mentioned that the measurement conditions in our channels do not match those of RT-DC due to the use of a different objective lens (20x instead of 40x), different carrier medium (higher methylcellulose concentration) and different stresses deforming the sample (extension vs. shear). Furthermore, our method operates at longer time-scales than RT-DC (~ 100 ms instead of ~ 5 ms). Thus, the RT-DC data are expected to merely provide a rough estimate.The maximum strain of validation sample B reaches values of 0.2 and higher for flow rates of 0.1 µL/s and 0.2 µL/s (see Fig. S3), which usually are assumed to be beyond the regime of linear elasticity. It is therefore surprising that the mechanical properties deviate very little over all four measured flow rates. This could indicate that the mechanical properties of validation sample B at flow rates ≥ 0.1 µL/s have to be interpreted with caution. The apparent viscosities in Fig. 4B show a different behavior for the two beads samples. While the sample validation A shows a decrease in viscosity, the values of sample validation B are roughly constant and lie in the expected range of 0.05 Pa s to 0.5 Pa s.^22,23^ The measured viscosity should be determined solely by the viscosity of the carrier medium, as the beads themselves are expected to behave almost purely elastic.^26^ The high viscosity values for validation sample A are not expected and could indicate the limitations of our method. Under the measurement conditions reported here, the signal-to-noise ratio of the strain could be too small, leading to inaccuracies during the fitting procedure of the mechanical model and, therefore, unrepresentative mechanical properties. Fig. 4C shows the relaxation time scale of the beads. Due to the linear relationship between this relaxation time τ, the Young’s modulus *E*, and the viscosity η, which is τ = η/*E*, it is clear that the relaxation times τ show the same trends as the viscosities η. Values on the order of 1 ms agree with previous studies and emphasize the almost purely elastic behavior of the beads.^26^ This is further supported by our unsuccessful attempt to fit the power-law rheology to the data. Although the algorithm converged to power-law exponents around zero, it resulted in large error values (see Fig. S4).

### Mechanical characterization of cells

To showcase the sensitivity to changes in the mechanical properties of biological samples, we measured HL60 cells with and without latB treatment. LatB is known to inhibit actin polymerization, therefore weakening the cytoskeleton, and softening cells.^24,35^We computed stress and strain depending on time instead of position as mentioned in the previous section. Mechanical models described in Eq. 8 and Eq. 12 were fitted to the strain and stress data. In contrast to the previous section, fitting included an additional parameter, ε_offset_, to account for the cells not being perfectly round. The results of fitting the Kelvin-Voigt model are presented in Fig. 5.

**Figure 5.**
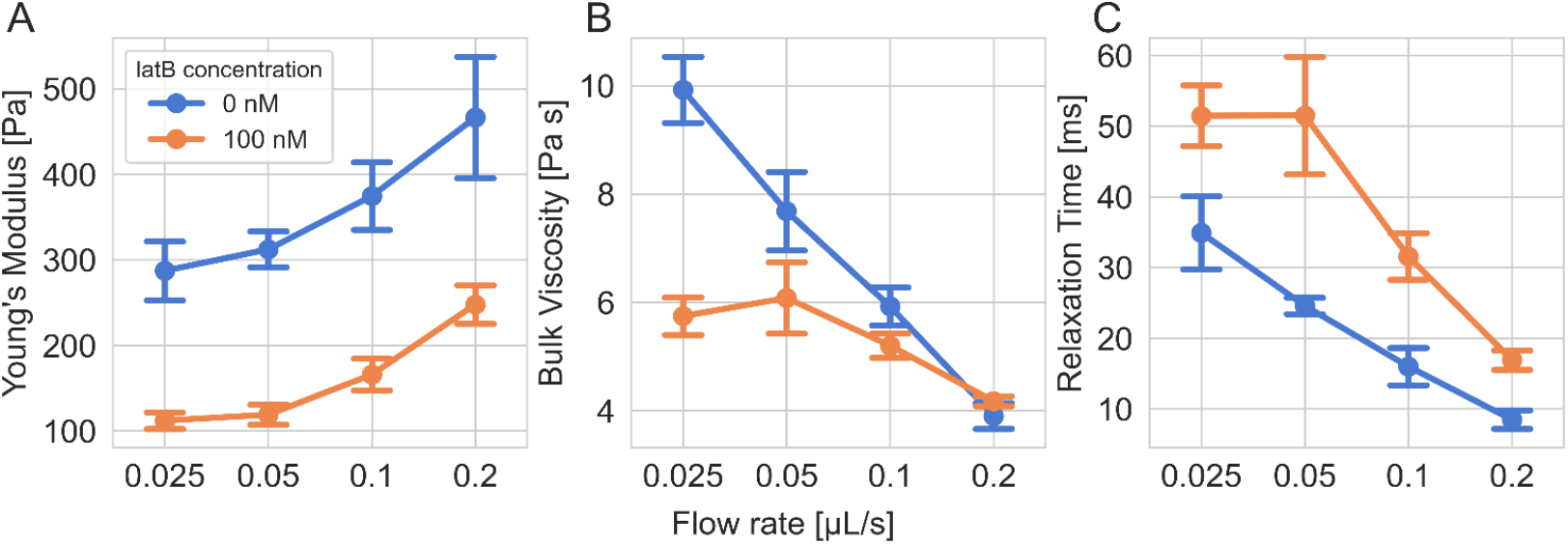
Mechanical characterization of HL60 cells treated with actin-modifying latrunculin B. A) Apparent Young’s modulus, *E*, of HL60 cells. B) Apparent viscosity, η. C) Relaxation time, τ. The data points represent mean values and standard deviations of three measurements.

The Young’s moduli shown in Fig. 5A show a clear difference between treated and untreated cells: LatB caused the cells to be 1.8-2.7 times softer than the untreated cells. The absolute values are comparable to earlier reports.^22,23^ Furthermore, both samples show Young’s moduli increasing with the flow rate. This apparent strain stiffening could be a non-linear strain response due to large strain values (see Fig. S5), therefore leading to higher estimated Young’s moduli.^21,23,36,37^ Another reason could be the time scale of deformation, as cell mechanical properties are generally time scale dependent.^38,39^ This would cause a different response to fast deformations at higher flow rates compared to slower deformations at lower flow rates, and was demonstrated for HL60 cells by Urbanska *et al*.^24^The apparent viscosity values we report here (Fig. 5B) are comparable to previous results.^22,23^ At the lowest flow rate, latB treated cells show a viscosity of about half the value of the untreated cells, while at the higher flow rates, the apparent viscosities are similar. With higher flow rates, the apparent viscosities of both samples decrease. This suggests shear thinning behavior and has been observed for HL60 cells before.^40^ Relaxation times are presented in Fig. 5C and show a decreasing trend with flow rate. The untreated sample has a shorter relaxation time, indicating an overall stiffer, more elastic response. The results of the power-law analysis are shown in Fig. 6. The material property *c*_β_, depicted in Fig. 6A, increases with flow rate, as does the power-law exponent β (Fig. 6B), which can be interpreted as a more liquid-like state. An apparent fluidization with increased external stress (= higher flow rate) has been observed before^41^ and was attributed to passive processes, such as alignment of fibers. LatB treated cells show a higher power-law exponent, which we attribute to a weakened actin cortex. Power-law exponents of β = 0.3 to β = 0.4 have been observed before for HL60 cells^20,22^, which is slightly higher compared to our observations of β = 0.22 ± 0.02 (untreated cells) and β = 0.34 ± 0.02 (latB treated cells). Overall, the power-law analysis indicates a rather elastic than viscous behavior of HL60 cells in our hyperbolic channels.

**Figure 6.**
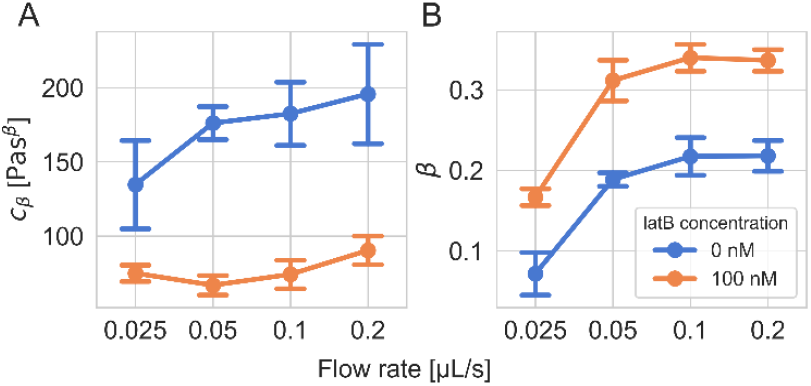
Power-law rheology of HL60 cells treated with latB. A) Material property *c*_β_. B) Power-law exponent over flow rate. The data points represent mean values and standard deviation of three measurements.

These results showcase the sensitivity of our method to differences in the viscoelastic properties of cells and drug induced changes of the cytoskeleton. The apparent discrepancy between the observed stiffening of the cells according to the Kelvin-Voigt model and the fluidization of the cells according to the power-law is worth mentioning, yet a direct comparison of these two different models is not meaningful; the Kelvin-Voigt model can be characterized by a single time scale while power-law rheology comprises a broad distribution of time scales.^38^ In general, a few points of improvement may be mentioned here. Using more sophisticated analysis methods, such as the one reported by Abuhattum *et al*.^42^, would allow to determine the complex shear modulus, *G*^∗^, and therefore provide a more detailed viscoelastic characterization. Furthermore, analysis is carried out on the combined bulk data, not on individual cells. This is due to the size of the measurement region which is currently not covered in a single field of view. By reducing the size of the measurement region, e.g. starting closer to the hyperbolic part and ending shortly after it ends, the overall acquisition time can be shortened. Furthermore, a camera with a larger field of view or an objective lens with lower magnification could be used to cover a larger part of the channel in a single measurement, although this would decrease resolution and cause larger uncertainties in the strain computations. Such development would enable single cell analysis, allowing to explore a broader spectrum of applications.

## Conclusions

In this study, we presented a microfluidic method for measuring the viscoelasticity of cells and cell-like objects using a hyperbolic channel. We demonstrated the combination of a hyperbolic channel design with 3D-hydrodynamic focusing in a single-layer soft lithography-based microfluidic chip. A channel height much larger than the size of cells and focusing of the cells in the center plane of the channel helped to keep shear stresses from the top and bottom of the channel minimal. We used PAAm microgel beads to characterize the stresses in the channel without having to characterize the rheology of the carrier medium due to their almost purely elastic behavior. We validated our approach using PAAm microgel beads and achieved similar results compared to other methods.^22,28^ Furthermore, we showed the sensitivity of our method to drug-induced changes of the cytoskeleton, demonstrating the potential as a universally applicable tool to investigate mechanical properties of cells. It could be used for a large-scale screening of drug libraries to identify substances that can alter the mechanical phenotype of a cell in yet unknown pathways. As our approach quantifies not only the Young’s moduli, but also the viscosities of cells, it has an advantage over other techniques, such as RT-DC, and could provide a deeper understanding of cell mechanics.^29,43^ Altogether, the method presented here is a promising approach for microfluidic-based viscoelastic characterization of biological cells.

## Author contributions

Benedikt Hartmann: Conceptualization, Data Curation, Formal analysis, Investigation, Methodology, Project administration, Software, Validation, Visualization, Writing – original draft, Writing – review & editing; Felix Reichel: Conceptualization, Methodology, Supervision, Writing – review & editing; Conrad Möckel: Formal analysis, Software, Writing – review & editing; Jochen Guck: Conceptualization, Funding acquisition, Project administration, Resources, Supervision, Writing – review & editing.

## Data availability

The raw data is available on the Deformability Cytometry Open Repository (DCOR): http://hdl.handle.net/21.11101/0000-0007-FE51-0.Python scripts used for analysis can be found at http://hdl.handle.net/21.11101/0000-0007-FE52-F.

## Acknowledgements

We want to acknowledge Shada Abuhattum and Marketa Kubankova for fruitful discussions on this study. We would like to thank Cornelia Liebers, Manuela Hauke and Christine Schweitzer for taking care of cell culture. We thank Ruchi Goswami, Parth Patel and Salvatore Girardo, part of the Core Facility Lab-on-a-Chip at Max-Planck-Zentrum für Physik und Medizin, for providing the PAAm microgel beads and the microfluidic chips used in this study. The authors acknowledge financial support by the Max Planck Society to JG.

