## Supplementary Material for "Viscoelastic characterization of cells in microfluidic channels with 3D hydrodynamic focusing"

### Supplementary materials and methods

Real-Time Deformability Cytometry

The Young's modulus and the size of the PAAm microgel beads used in this work were determined using real-time deformability cytometry (RT-DC)[1]. Experiments were carried out on an AcCellerator (Zellmechanik Dresden, Germany). All bead samples were measured in a channel with 30 µm x 30 µm cross section at several different flow rates. The carrier medium was 0.6 %w/v methylcellulose in phosphate buffered saline. We use the model introduced by Wittwer et al.[2] and the viscosity model described by Büyükurganci et al.[3] to calculate the Young's modulus.

### Supplementary tables

Table S1: PAAm microgel beads: Monomer concentration

| Sample | Calibration A | Calibration B | Calibration C | Validation A | Validation B |
|---|---|---|---|---|---|
| **Total monomer concentration [%w/v]** | 6.0 | 4.5 | 5.2 | 5.6 | 4.5 |

Table S2: PAAm microgel beads: Fit parameters of Kelvin-Voigt fits

| Sample | Flow rate [µl/s] | E [Pa] | ΔE [Pa] | η [Pa s] | $\Delta\eta$ [Pa s] | τ [s] | Δτ [s] | $R^2$ |
|---|---|---|---|---|---|---|---|---|
| Validation A | 0.025 | 1340 | 18 | 5.639 | 1.650 | 0.0042 | 0.0012 | 0.9445 |
| | 0.05 | 1477 | 15 | 3.107 | 0.706 | 0.0021 | 0.0005 | 0.9695 |
| | 0.1 | 1423 | 12 | 1.150 | 0.264 | 0.0008 | 0.0002 | 0.9817 |
| | 0.2 | 1357 | 10 | 0.678 | 0.149 | 0.0005 | 0.0001 | 0.9852 |
| Validation B | 0.025 | 523 | 13 | 0.596 | 0.041 | 0.0011 | 0.0001 | 0.9649 |
| | 0.05 | 406 | 4 | 0.238 | 0.008 | 0.0006 | 0.0000 | 0.9921 |
| | 0.1 | 434 | 2 | 0.232 | 0.012 | 0.0005 | 0.0000 | 0.9966 |
| | 0.2 | 406 | 1 | 0.263 | 0.022 | 0.0006 | 0.0001 | 0.9964 |

Table S3: PAAm microgel beads: Fit parameters of Power-law fits

| Sample | Flow_rate [µl/s] | $c_\beta$ [Pa $s^\beta$] | $\Delta c_\beta$ [Pa $s^\beta$] | β | Δβ | $R^2$ |
|---|---|---|---|---|---|---|
| Validation A | 0.025 | 41.522 | 138.806 | 0.005 | 0.016 | 0.9695 |
| | 0.05 | 176.538 | 128.098 | 0.020 | 0.016 | 0.9644 |
| | 0.1 | 1.653 | 147.346 | 0.000 | 0.015 | 0.9631 |
| | 0.2 | 1.574 | 135.704 | 0.000 | 0.015 | 0.9671 |
| Validation B | 0.025 | 0.434 | 173.967 | 0.000 | 0.043 | 0.8126 |
| | 0.05 | 0.383 | 91.315 | 0.000 | 0.028 | 0.8897 |
| | 0.1 | 0.419 | 62.583 | 0.000 | 0.021 | 0.9348 |
| | 0.2 | 0.404 | 49.114 | 0.000 | 0.018 | 0.9534 |

Table S4: HL60 cells: Fit parameters of Kelvin-Voigt fits

| Sample | Flow rate [µl/s] | Rep. | E [Pa] | ΔE [Pa] | η [Pa s] | $\Delta\eta$ [Pa s] | τ [s] | $\Delta\tau$ [$s$] | $\varepsilon_{offset}$ | $\Delta\varepsilon_{offset}$ | $R^2$ |
|---|---|---|---|---|---|---|---|---|---|---|---|
| 0 nM latB | 0.025 | 1 | 326 | 6 | 9.93 | 0.47 | 0.0305 | 0.0015 | 0.0468 | 0.0007 | 0.9941 |
| | | 2 | 259 | 4 | 10.53 | 0.37 | 0.0406 | 0.0015 | 0.0437 | 0.0007 | 0.9949 |
| | | 3 | 276 | 5 | 9.31 | 0.45 | 0.0337 | 0.0017 | 0.0455 | 0.0008 | 0.9930 |
| | 0.05 | 1 | 314 | 4 | 8.12 | 0.19 | 0.0259 | 0.0007 | 0.0572 | 0.0008 | 0.9965 |
| | | 2 | 332 | 5 | 8.09 | 0.26 | 0.0243 | 0.0009 | 0.0642 | 0.0009 | 0.9951 |
| | | 3 | 291 | 4 | 6.84 | 0.20 | 0.0236 | 0.0007 | 0.0629 | 0.0008 | 0.9958 |
| | 0.1 | 1 | 410 | 6 | 5.54 | 0.16 | 0.0135 | 0.0004 | 0.0755 | 0.0013 | 0.9945 |
| | | 2 | 381 | 4 | 5.99 | 0.12 | 0.0157 | 0.0004 | 0.0780 | 0.0010 | 0.9971 |
| | | 3 | 332 | 5 | 6.23 | 0.16 | 0.0187 | 0.0006 | 0.0876 | 0.0015 | 0.9949 |
| | 0.2 | 1 | 519 | 8 | 3.74 | 0.11 | 0.0072 | 0.0002 | 0.0895 | 0.0018 | 0.9942 |
| | | 2 | 495 | 9 | 4.17 | 0.13 | 0.0084 | 0.0003 | 0.0978 | 0.0019 | 0.9939 |
| | | 3 | 386 | 8 | 3.79 | 0.12 | 0.0098 | 0.0004 | 0.1032 | 0.0028 | 0.9912 |
| 100 nM latB | 0.025 | 1 | 115 | 1 | 5.45 | 0.11 | 0.0473 | 0.0010 | 0.0854 | 0.0011 | 0.9977 |
| | | 2 | 120 | 1 | 6.13 | 0.12 | 0.0512 | 0.0011 | 0.0942 | 0.0010 | 0.9981 |
| | | 3 | 101 | 1 | 5.66 | 0.12 | 0.0559 | 0.0013 | 0.0964 | 0.0013 | 0.9973 |
| | 0.05 | 1 | 127 | 2 | 5.38 | 0.13 | 0.0422 | 0.0012 | 0.1187 | 0.0023 | 0.9946 |
| | | 2 | 124 | 2 | 6.70 | 0.14 | 0.0540 | 0.0014 | 0.1209 | 0.0021 | 0.9957 |
| | | 3 | 106 | 2 | 6.15 | 0.14 | 0.0583 | 0.0017 | 0.1230 | 0.0028 | 0.9938 |
| | 0.1 | 1 | 170 | 4 | 5.00 | 0.13 | 0.0295 | 0.0010 | 0.1400 | 0.0038 | 0.9919 |
| | | 2 | 182 | 3 | 5.44 | 0.11 | 0.0298 | 0.0008 | 0.1613 | 0.0026 | 0.9959 |
| | | 3 | 146 | 3 | 5.15 | 0.14 | 0.0354 | 0.0013 | 0.1567 | 0.0044 | 0.9913 |
| | 0.2 | 1 | 255 | 5 | 4.12 | 0.11 | 0.0162 | 0.0005 | 0.1625 | 0.0042 | 0.9931 |
| | | 2 | 266 | 6 | 4.28 | 0.12 | 0.0161 | 0.0006 | 0.1956 | 0.0040 | 0.9938 |
| | | 3 | 223 | 6 | 4.12 | 0.13 | 0.0185 | 0.0007 | 0.1956 | 0.0054 | 0.9911 |

Table S5: HL60 cells: Fit parameters of Power-law fits

| Sample | Flow rate [µl/s] | Rep. | $c_\beta$ [Pa $s^\beta$] | $\Delta c_\beta$ [Pa $s^\beta$] | β | Δβ | $\varepsilon_{offset}$ | $\Delta\varepsilon_{offset}$ | $R^2$ |
|---|---|---|---|---|---|---|---|---|---|
| 0 nM latB | 0.025 | 1 | 154.860 | 20.656 | 0.070 | 0.016 | 0.0428 | 0.0012 | 0.9882 |
| | | 2 | 148.705 | 8.685 | 0.099 | 0.013 | 0.0383 | 0.0013 | 0.9890 |
| | | 3 | 100.348 | 26.020 | 0.046 | 0.017 | 0.0417 | 0.0013 | 0.9872 |
| | 0.05 | 1 | 178.920 | 2.762 | 0.194 | 0.008 | 0.0449 | 0.0012 | 0.9945 |
| | | 2 | 185.486 | 2.729 | 0.193 | 0.007 | 0.0507 | 0.0011 | 0.9955 |
| | | 3 | 163.861 | 2.261 | 0.179 | 0.007 | 0.0490 | 0.0011 | 0.9951 |
| | 0.1 | 1 | 197.841 | 2.778 | 0.201 | 0.006 | 0.0547 | 0.0012 | 0.9966 |
| | | 2 | 191.205 | 3.207 | 0.207 | 0.007 | 0.0600 | 0.0014 | 0.9956 |
| | | 3 | 158.222 | 2.019 | 0.244 | 0.004 | 0.0648 | 0.0012 | 0.9978 |
| | 0.2 | 1 | 221.063 | 2.581 | 0.200 | 0.004 | 0.0632 | 0.0012 | 0.9983 |
| | | 2 | 208.358 | 2.243 | 0.216 | 0.004 | 0.0716 | 0.0010 | 0.9988 |
| | | 3 | 157.760 | 1.753 | 0.238 | 0.003 | 0.0702 | 0.0013 | 0.9986 |
| 100 nM latB | 0.025 | 1 | 75.555 | 1.062 | 0.165 | 0.007 | 0.0670 | 0.0019 | 0.9950 |
| | | 2 | 80.074 | 1.453 | 0.157 | 0.009 | 0.0799 | 0.0021 | 0.9939 |
| | | 3 | 69.166 | 1.081 | 0.178 | 0.008 | 0.0782 | 0.0023 | 0.9938 |
| | 0.05 | 1 | 70.696 | 0.832 | 0.284 | 0.005 | 0.0904 | 0.0017 | 0.9977 |
| | | 2 | 70.601 | 0.868 | 0.318 | 0.005 | 0.0981 | 0.0016 | 0.9979 |
| | | 3 | 59.179 | 0.551 | 0.333 | 0.004 | 0.0950 | 0.0014 | 0.9986 |
| | 0.1 | 1 | 74.936 | 1.094 | 0.334 | 0.005 | 0.1060 | 0.0023 | 0.9976 |
| | | 2 | 83.311 | 1.150 | 0.327 | 0.004 | 0.1331 | 0.0019 | 0.9983 |
| | | 3 | 64.258 | 0.838 | 0.359 | 0.004 | 0.1212 | 0.0022 | 0.9981 |
| | 0.2 | 1 | 91.530 | 1.160 | 0.338 | 0.004 | 0.1275 | 0.0019 | 0.9987 |
| | | 2 | 99.245 | 1.261 | 0.322 | 0.004 | 0.1611 | 0.0018 | 0.9990 |
| | | 3 | 80.133 | 1.352 | 0.349 | 0.005 | 0.1567 | 0.0028 | 0.9979 |

Fig. S1: PAAm microgel beads: Size measured by RT-DC

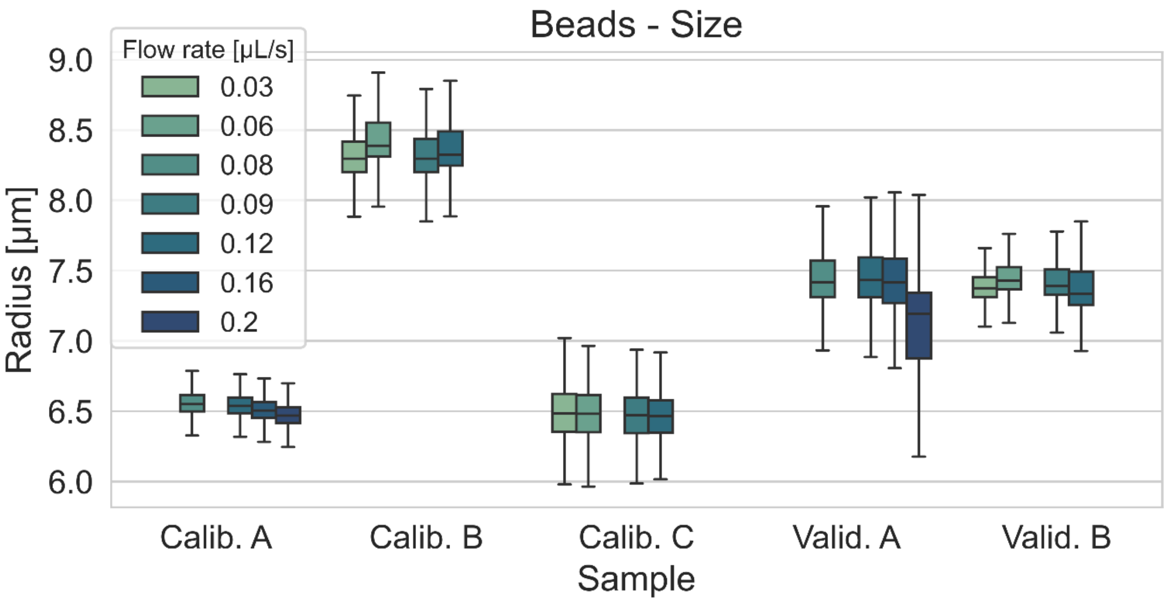


Fig. S2: PAAm microgel beads: Young's modulus measured by RT-DC

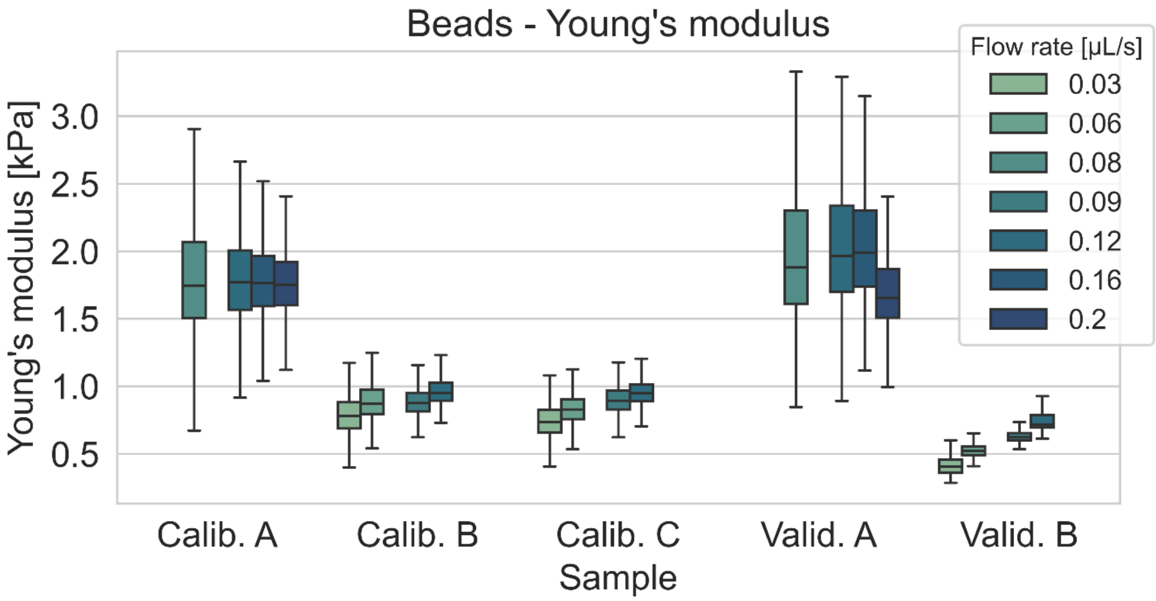

Fig. S3: PAAm microgel beads: Example plot for fits of mechanical models to sample validation B measured at 0.1 mL/s

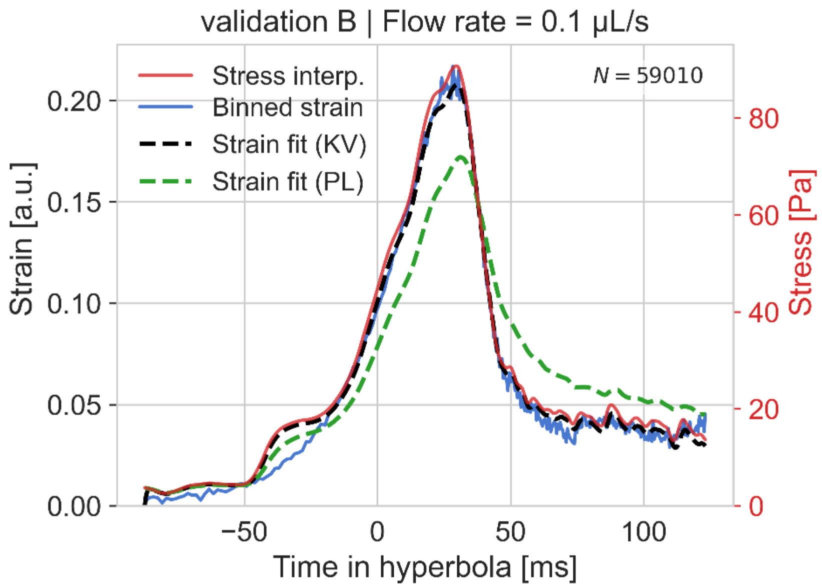


Fig. S4: PAAm microgel beads: Power-law rheology

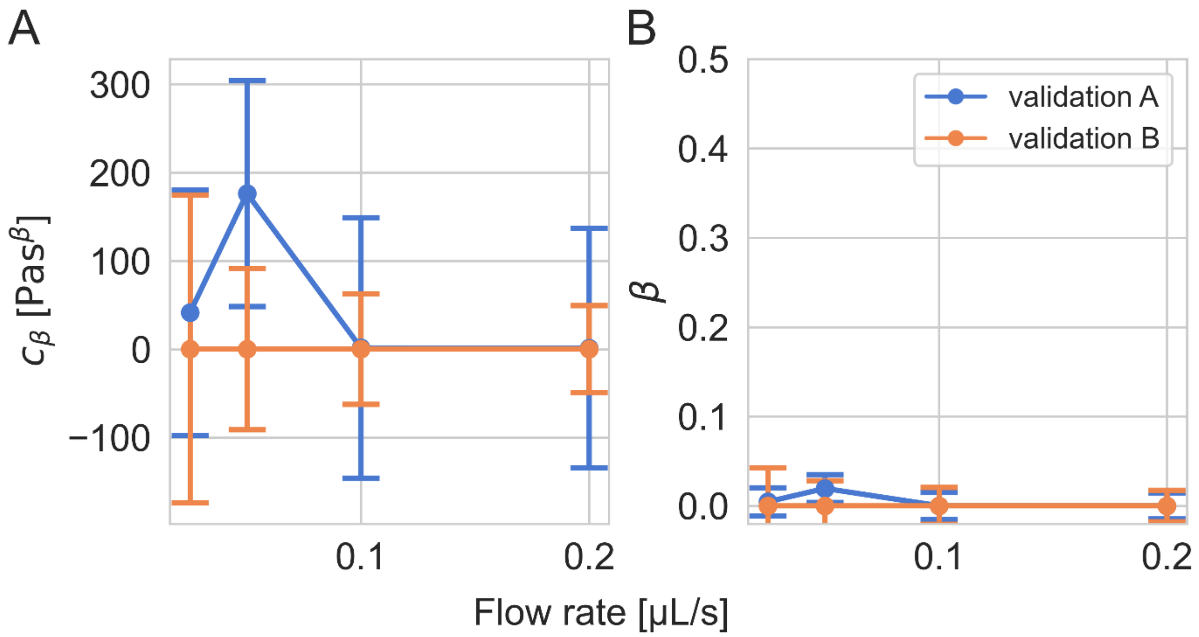

Fig. S5: HL60 cells: Example plots for fits of mechanical models. A) Sample without latB treatment. B) Sample treated with 100 nM latB.

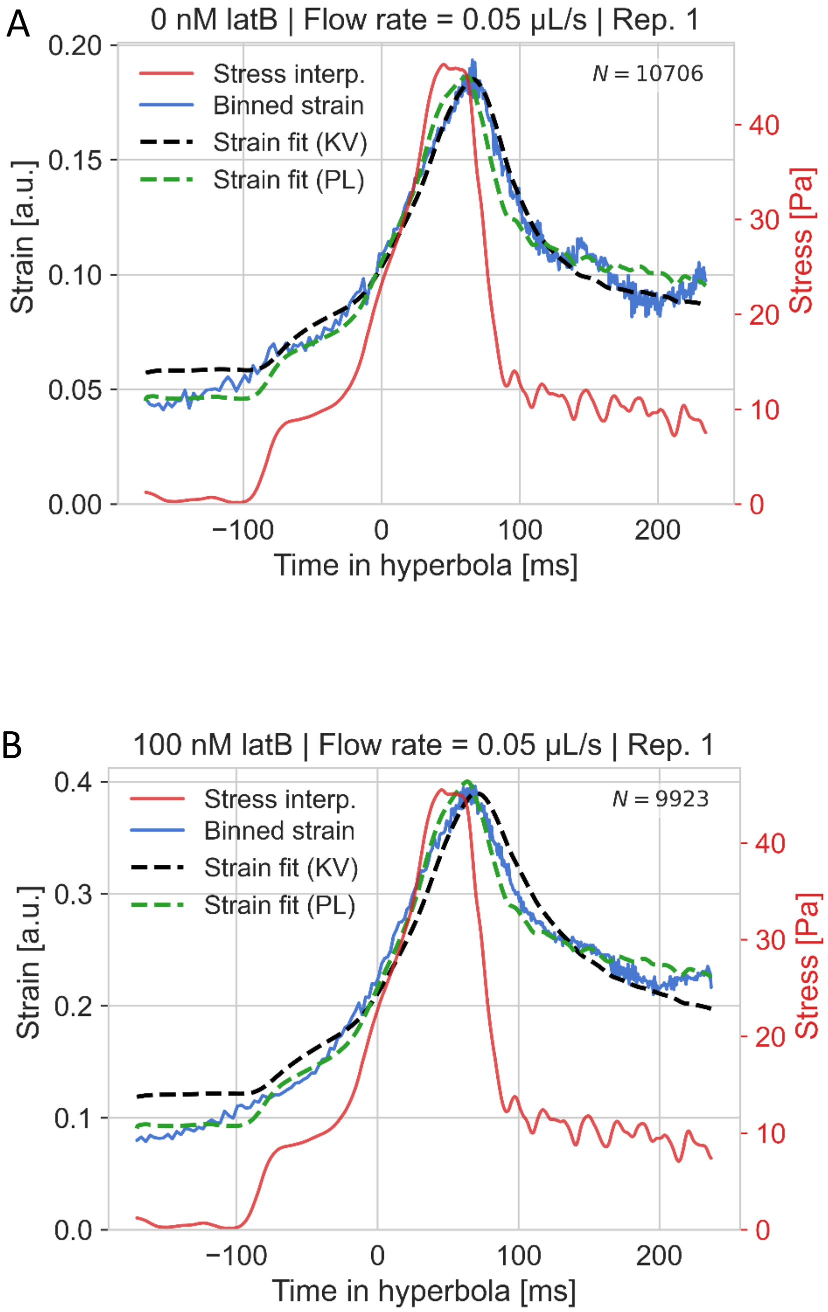
